# Subgroup-specific gene expression profiles and mixed epistasis in chronic lymphocytic leukemia

**DOI:** 10.1101/2021.04.16.440134

**Authors:** Almut Lütge, Junyan Lu, Jennifer Hüllein, Tatjana Walther, Leopold Sellner, Bian Wu, Richard Rosenquist, Christopher C. Oakes, Sascha Dietrich, Wolfgang Huber, Thorsten Zenz

**Affiliations:** Genome Biology Unit, EMBL, Heidelberg, 69117, Germany; Department of Molecular Life Sciences, University of Zurich, Zurich, Switzerland; SIB Swiss Institute of Bioinformatics, University of Zurich, Zurich, Switzerland; Molecular Therapy in Hematology and Oncology & Department of Translational Oncology, NCT and DKFZ, Heidelberg, Germany; Department of Medicine V, Heidelberg University Hospital, Heidelberg, Germany; Cancer Center, Union Hospital, Tongji Medical College, Huazhong University of Science and Technology, Wuhan 430022, China; Department of Molecular Medicine and Surgery, Karolinska Institutet, Stockholm, Sweden; Clinical Genetics, Karolinska University Hospital, Solna, Sweden; Department of Internal Medicine, Division of Hematology, The Ohio State University, Columbus, USA; Department of Medical Oncology and Hematology, University Hospital Zurich, Zurich, Switzerland

## Abstract

Despite the extensive catalogue of recurrent mutations in chronic lymphocytic leukaemia (CLL), the diverse molecular driving events and the resulting range of disease phenotypes remain incompletely understood. To study the molecular heterogeneity of CLL, we performed RNA-sequencing on 184 CLL patient samples. Unsupervised analysis revealed two major independent axes of gene expression variation: the first one aligned with the mutational status of the immunoglobulin heavy variable (IGHV) genes, and concomitantly, with the three-group stratification of CLL by global DNA methylation pattern, and affected biological functions including B- and T-cell receptor signaling. The second one aligned with trisomy 12 status and affected chemokine signaling. Furthermore, we searched for differentially expressed genes associated with gene mutations and copy-number aberrations and detected strong signatures for *TP53, BRAF* and *SF3B1*, as well as for del(11)(q22.3), del(17)(p13) and del(13)(q14) beyond the dosage effect. We discovered strong non-additive effects (i.e., genetic interactions, or epistasis) of IGHV mutation status and trisomy 12 on multiple phenotypes, including the expression of 893 genes. Multiple types of epistasis were observed, including synergy, buffering, suppression and inversion. Our study reveals previously underappreciated gene expression signatures for (epi)genomic variants in CLL and the presence of epistasis between them. The findings will serve as a reference for a functional resolution of CLL molecular heterogeneity.

## Introduction

Chronic lymphocytic leukemia (CLL) etiology has been linked to abnormal B-cell receptor (BcR) activation and gene mutations targeting multiple pathways, including DNA damage pathways (*TP53, ATM*), *NOTCH* signaling (*NOTCH1, FBXW7, MED12*)^1–4^ and the spliceosome (*SF3B1*)^5,6^. In addition, the IGHV mutation status, the result of a physiological mutation and maturation process, reflects the tumor’s cell type of origin and is one of the strongest predictors of clinical behavior ^7^. Several genetic subgroups of CLL are known that show profound differences in clinical course, presentation and outcome^8,9^, although considerable variability remains within subgroups.

Gene expression profiling has the potential to provide a better understanding of the functional role of mutations and may help to dissect the disease heterogeneity. Indeed, previous studies of CLL transcriptomes have found substantial variability^10,11^, however, it has been a surprise how little of that variability could be associated with the genetic subgroups or other properties of the disease. For instance, Ferreira et al.^10,11^ found only a few robust gene expression changes associated with the major cancer driver mutations of CLL. IGHV mutation status only accounted for 1.5% of the overall variance in their study. The study reported two subgroups, termed C1/C2, as a predictor of clinical outcome independent of known molecular disease groups. A later reanalysis of the data suggested a relation of C1/C2 to sample processing^12^.

Overall, the relations between prominent genetic events that have significant impact on disease course and the gene expression programmes of CLL have remained unclear. Among possible explanations for this scarcity of associations are small sample sizes, confounding effects of multiple cytogenetic abnormalities or confounding technical variations. More recent studies have thus collected larger cohorts with focus on a particular genetic aberration. Abruzzo et al.^13^ identified a set of dysregulated and potentially targetable pathways in CLL with trisomy 12. Herling et al.^14^ developed a 17-gene signature that can identify a subset of treatment-naive patients with IGHV-unmutated CLL (U-CLL) who might substantially benefit from treatment with FCR (fludarabine, cyclophosphamide and rituximab) chemoimmunotherapy. These findings underline the importance of transcriptional changes in CLL.

Here, we profiled 184 CLL samples using RNA-sequencing. After careful control of technical variations, and of possible confounding between genetic variants, we searched for transcriptomic signatures and pathway activity changes associated with the major recurrent genetic alternations in CLL. Furthermore, as a step towards gaining a better understanding of functional interdependencies between mutations in a tumor, we used a quantitative model of genetic interactions to identify non-additive effects of mutations on gene expression profiles.

## Material and Methods

### Data acquisition

#### RNA-sequencing

We selected 184 CLL patient samples for RNA-sequencing. Patients were recruited between 2011-2017 with informed consent. The population was representative for a tertiary referral center. The majority of patients (177 out of 184) showed a typical CLL phenotype and 5 patients were diagnosed with atypical CLL. In total 92 patients had undergone some kind of medical treatment. All patient characteristics are shown in Supplemental Table S1. RNA isolation and library preparation were performed as described before^15^. In short, total RNA was isolated from patient blood samples (CD19+ presorted n=161) using the RNA RNeasy mini kit (Qiagen). RNA quantification was performed with a Qubit 2.0 Fluorometer. RNA integrity was evaluated with an Agilent 2100 Bioanalyzer, and samples with RNA integrity number (RIN) <8 were excluded. Sequencing libraries were prepared according to the Illumina TruSeq RNA sample preparation v2 protocol. Samples were paired-end sequenced at the DKFZ Genomics and Proteomics Core Facility. Two to three samples were multiplexed per lane on Illumina HiSeq 2000, Illumina HiSeq3000/4000 or Illumina HiSeqX machines.

Raw RNA-sequencing reads were demultiplexed, and quality control was performed using FastQC^16^ version 0.11.5. STAR^17^ version 2.5.2a was used to remove adapter sequences and map the reads to the Ensembl human reference genome release 75 (Homo sapiens GRCh37.75). STAR was run in default mode with internal adapter trimming using the *clip3pAdapterSeq* option. Mapped reads were summarized into per gene counts using htseq-count^18^ version 0.9.0 with default parameters and union mode. Thus, only reads unambiguously mapping to a single gene were counted. The count data were imported into R^19^ (version 3.6) for subsequent analysis.

#### Somatic variants

Mutation calls for 66 distinct gene mutations and 22 structural variants had been generated in a previous study for 143 out of 184 CLL samples through targeted sequencing, whole-exome sequencing and whole-genome sequencing^15^. For the remaining 41 samples, we generated additional targeted and whole-genome sequencing data and called variants using the same pipeline. Statistical analyses were restricted to variants found in at least 5 patients, i.e., to 14 gene mutations (*BRAF, NOTCH1, SF3B1, TP53, KRAS, ATM, MED12, EGR2, KLHL6, ACTN2, MGA, NFKBIE, PCLO, XPO1*), and 9 copy-number aberrations (CNAs): (trisomy 12, del(11)(q22.3), del(13)(q14), del(17)(p13), del(8)(p12), gain(8)(q24), gain(2)(p25.3), del(15)(q15.1), gain(14)(q32)) (Supplemental Figure S2B). In addition, the somatic hypermutation status of the immunoglobulin heavy variable (IGHV) and a CLL subtype classification defined by global patterns of CpG methylation level^20,21^ were recorded. In this paper, we discuss results for variants for which at least 100 genes were detected as differentially expressed at a false discovery rate (FDR) of 0.01 according to the method of Benjamini-Hochberg^22^: 4 cCNAs, 3 gene mutations and IGHV mutation status.

### Statistical analysis

#### Exploratory data analysis: PCA and clustering

Statistical analyses were performed using R^19^ version 3.6. The exploratory data analysis was performed on data normalized and transformed using the variance stabilizing transformation (VST) provided by the DESeq2 package^23^. The 500 most variable genes were used in the principal component analysis (PCA) and hierarchical clustering. PCA was done using the *prcomp* function with *scale*. Hierarchical clustering with the *ward*.*D2* method was performed on sample Euclidean distances computed on the scaled gene expression values. The *complexHeatmaps* package^24^ was used to visualize results.

#### Batch effect estimation

Transcriptome data were generated over a period of four years and platforms were changed with technological development during the period of sequencing, which led to changes in sequencing depth and read length (101, 125 and 151 nucleotides). Therefore, we considered the possibility of batch effects in the data due to platform differences^25^. Before adapter trimming we found a higher fraction of reads that contained adapter sequences in batches with longer reads. These resulted in batch dependent mapping to pseudogenes. After adapter trimming we did not detect differences in mapping towards pseudogenes or any associations between the top 10 principal components or the investigated genetic variants and different batches (Supplemental Figure S1).

### Differential expression analysis

For each of the 23 genetic alterations (13 gene mutations, 10 CNAs) and the IGHV mutation status, differentially expressed genes were identified using the Gamma-Poisson generalized linear modeling (GLM) approach of DESeq2, version 1.16.1^23,26^. Because of the large effects of IGHV mutation status and trisomy 12 on gene expression (as seen in the exploratory data analysis), these two variables were used as blocking factors in the models for each of the 22 remaining variants. In the model for IGHV mutation status, trisomy 12 was used as a blocking factor, and vice versa. Genetic interactions were identified by testing for an interaction term between the two variables IGHV mutation status and trisomy 12. Separately in each of these 25 DESeq2 analyses, the method of Benjamini and Hochberg^22^ was applied to account for multiple testing and control FDR of 0.01.

### Gene set enrichment analysis

Gene set enrichment analysis^27^ was performed using the R package clusterProfiler^28^ version 3.12.0 based on ranked gene statistics from DESeq2. Hallmark and KEGG gene set collections version 4.0 were downloaded from MSigDB^29^. The significance of gene sets was determined using a permutation null (B=1000). P-values were adjusted for multiple testing using the method of Benjamini and Hochberg^22^.

## Data Sharing Statement

RNA-sequencing data are available at European Genome-phenomeArchive (EGA) under accession number EGAS00001001746. All code to reproduce this analysis is available at https://github.com/almutlue/transcriptome_cll (DOI:10.5281/zenodo.4600322). Analysis code and outputs are deployed as a browsable workflowr^30^ site: https://almutlue.github.io/transcriptome_cll/index.html.

## Results

### Unsupervised analysis reveals major drivers of gene expression variability

We generated RNA-sequencing data from 184 CLL patient samples. To obtain a first overview of patterns of gene expression variability in CLL, we performed an unsupervised clustering analysis based on the 500 most variable genes (Figure 1A). This analysis showed a separation of distinct subgroups that largely coincided with IGHV mutation status/methylation epitype and the presence of trisomy 12. The role of IGHV mutation status and trisomy 12 was also reflected in the number of differentially expressed (DE) genes (>3000 DE genes) (Figure 1B). A similar separation was seen in a principal component analysis (PCA) (Figure 1C,D). The first principal component, which represented 11% of the variance, was associated with IGHV mutation status, while the second component separated samples based on trisomy 12. These results indicate that these two genetic variables shape gene expression in CLL to a previously unappreciated extent. We also considered a classification of CLL based on global DNA methylation levels into three groups^20,21^, a refinement of the binary grouping by IGHV mutation status (Figure 1E). Accordingly, the first principal component arranged the DNA methylation subgroups into the order low, intermediate and high programmed (LP, IP, HP). These results indicate that even though the three groups classification was discovered using DNA methylation data, it is equally apparent at the level of gene expression. Indeed, the global gene expression patterns shown in Figure 1 imply a further refinement into five major groups, namely LP, IP and HP each with and without trisomy 12.

**Figure 1.**
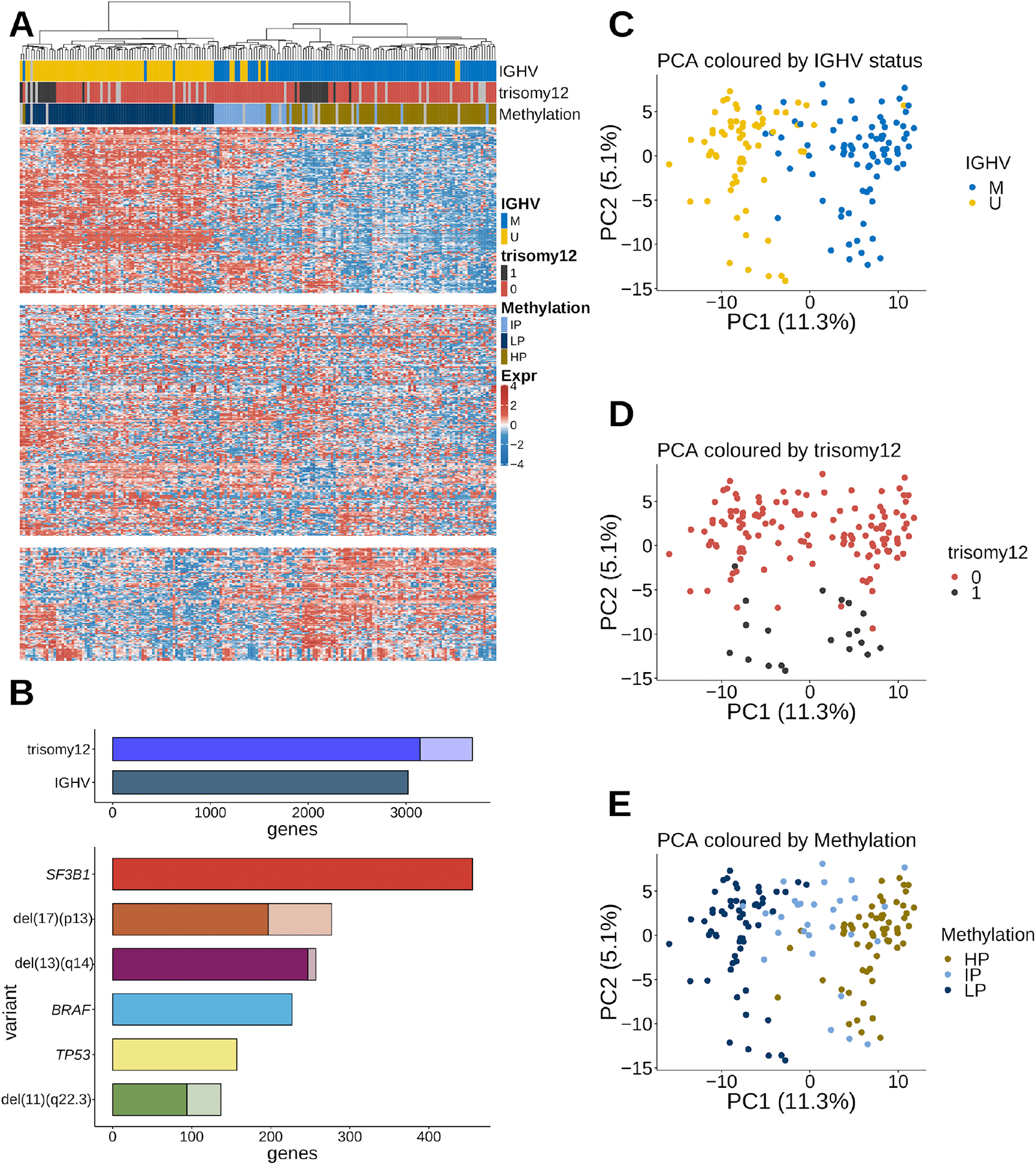
Gene expression variability in CLL: A) Heatmap of the gene expression counts. Samples (columns) are ordered in agreement with the hierarchical clustering based on the 500 most variable genes. Gene (rows) counts are row-centered log transformed (base 2). IGHV mutation status, methylation subgroups and trisomy 12 align with the clustering result. B) Number of differentially expressed genes (adjusted P-values < 0.01) for genomic markers in CLL. Lighter colors indicate genes located on the same chromosome as the respective genetic lesion (potential dosage effects). C) IGHV status is associated with the first principal component, which explains 11.3% of the variance. D) Trisomy 12 is associated with the second principal component, which explains 5.1% of the variance. E) Methylation subgroups split up along principal component 1.

The results of our unsupervised clustering analysis differ from those of a previous gene expression study, which also used unsupervised clustering of RNA-sequencing data to find novel subgroups of CLL termed C1/C2, marked by 600 differentially expressed genes and associated with BcR activation and outcome^10^. In our data, hierarchical clustering of the samples based on the measurements of these 600 genes only, indeed resulted in two main clusters. However, most of these genes showed low variability across samples and only 26 of them were among the 500 most variable genes (Supplemental Figure S2).

### Mutations modulate gene expression in CLL

We performed differential expression analysis to explore the effect of 23 recurrent genetic aberrations and the IGHV mutations status (Supplemental Figure S3). In total, we found 6 additional variants (besides trisomy 12 and the IGHV status) associated each with more than 100 differentially expressed genes. These were del(13)(q14), del(17)(p13), del(11)(q22), and *SF3B1, TP53* and *BRAF* mutations (Figure 1B). Complete tables are provided in the computational analysis transcript.

We compared previous findings from the literature for single genetic aberrations with differentially expressed genes in this study. Mutations in the splicing factor *SF3B1* gene showed more than 400 associations. Gene sets enriched in CLL with *SF3B1* mutations included “Cytokine-cytokine receptor interaction” and “Phosphatidylinositol signaling system” (Supplemental Figure S4A). Among differentially expressed genes, we found the chaperone gene *UQCC1* (Supplement Figure S4B), which has already been linked to *SF3B1* mutations by differential isoform usage^31^. We also found *PSD2, SRRM5* and *TNXB* (Supplemental Figure S3C-E) associated with *SF3B1* mutation. *TP53* is another commonly mutated gene in CLL and associated with inferior survival^1^. Differentially expressed genes in samples with *TP53* mutation were enriched in “Oxidative phosphorylation” and “p53 signaling pathway” (Supplement Figure S5A). The transcriptional regulator *CDK12* is upregulated in *TP53* mutated samples (Supplement Figure S4B). Further associations with *TP53* include *PGBD2, HYPK* and the p53 antagonist *MDM2*^*32*^ (Supplement Figure S4C-E).

### IGHV mutation status is linked to distinct gene expression changes

The highest number of differentially expressed genes was found in the comparison between IGHV-mutated (M-CLL) and U-CLL: 3275 genes. This result is in agreement with the PCA of Figure 1C and shows that IGHV mutation status is the main determinant of gene expression variability in CLL. It implies a much larger impact of IGHV status on the transcriptome than previously detected (11.3% instead of 1.5% ^10^) and is in line with the key impact of IGHV mutation status on clinical course and biology of disease ^7–9^. Genes previously found to be markers related to IGHV mutation status, including *CD38, LPL, ZAP70, SEPT10, ADAM29* and *PEG10*^*33–35*^, were also detected in our analysis (Supplemental Table 1).

To understand the pathways involved in U-CLL and M-CLL, we performed gene set enrichment analysis. Differentially expressed genes between IGHV groups were enriched in BcR signaling, T-cell receptor signaling and chemokine signaling pathways (Figure 2A). Within the BcR signaling gene set, we identified cell surface molecules (*CD19, CD22, CD81*) and NFAT and NF-κB to be downregulated in U-CLL. From the “T cell receptor signaling” gene set, *ZAP70, PAK* and p38 were upregulated in U-CLL, while *IL10* and *SHP1* were downregulated. Within chemokine signaling pathways, we found downregulation of *CXCR3* and *CXCR4* in U-CLL, while a set of interferons (*IFNB1, INFA21*) were upregulated.

**Figure 2.**
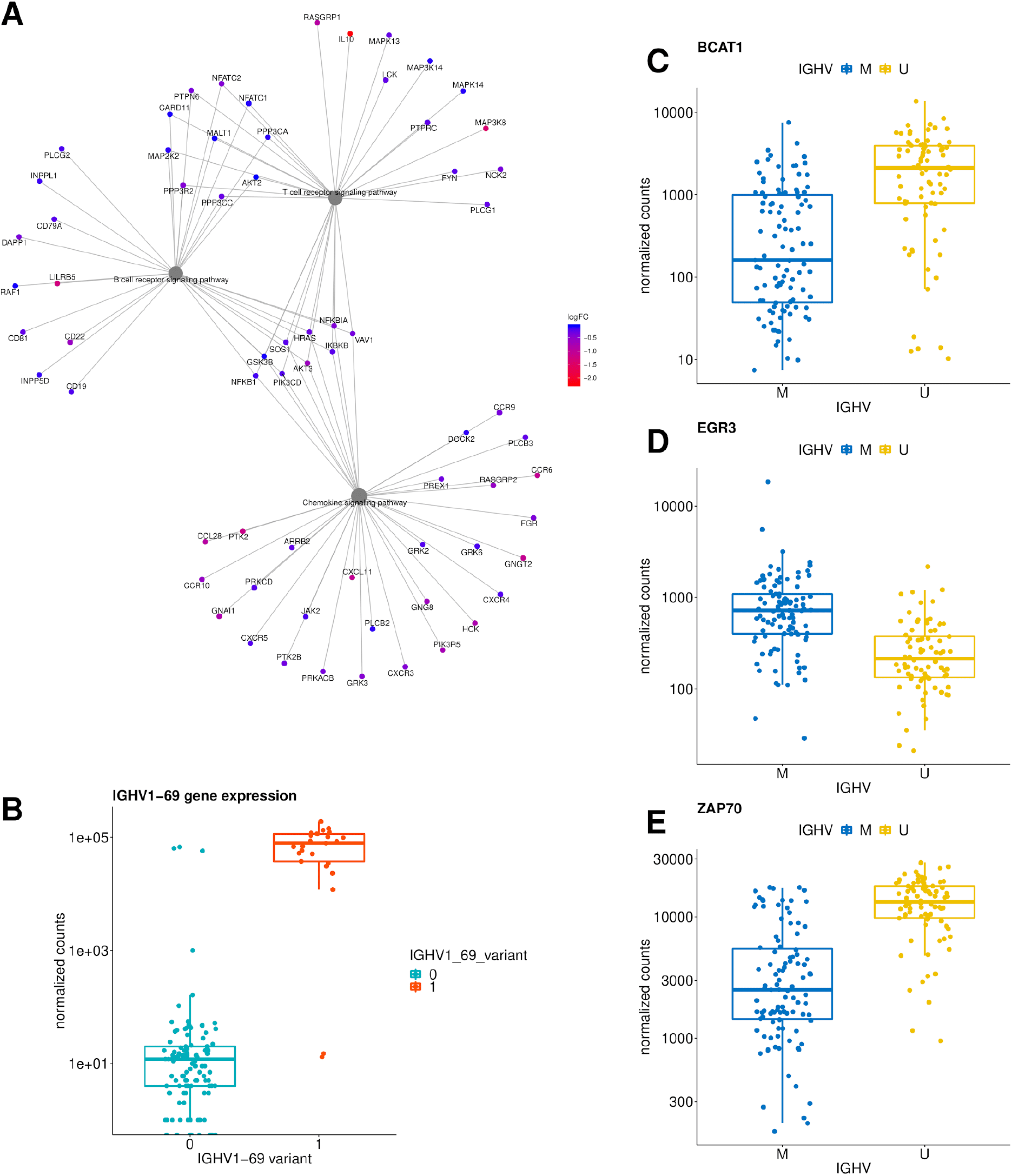
Gene expression changes between IGHV subgroups: A) Differentially expressed genes in enriched KEGG pathways for IGHV. B) IGHV1-69 expression by corresponding IGHV1-69 gene usage determined by IG gene analysis. C-E) Normalized gene counts for *BCAT1, EGR3* and *ZAP70* separated by IGHV mutation status.

IGHV genes were also found among the most differentially expressed genes, but showed heterogeneous expression within the U-CLL and M-CLL groups. As expected, commonly used IGHV genes (IGHV1-69 or IGHV4-34) were associated with U-CLL and M-CLL, respectively. Gene expression showed a strong relation to IG gene usage and it’s variant’s expression (Figure 2B). These data show that RNA-sequencing can be used to assess IG gene usage. Further genes associated with IGHV groups were *BCAT1, EGR3* and *ZAP70* (Figure 2C-E)

In summary, our data are in line with the major biological role of IGHV mutation status in CLL and provide a resource to identify deregulated pathways in the disease.

### Intermediate programmed methylation group forms an independent gene expression cluster

Based on the global DNA methylation pattern, the stratification of CLL by IGHV mutation status has recently been refined by introducing a categorization into LP, IP, and HP programmed samples, which are thought to represent the cell of origin ^20,21^. Based on gene expression data, we found these three groups along the first principle component. The IP group was placed between the LP and HP groups (Figure 1E). Thus, even though the groups were discovered on the basis of DNA methylation, a strong separation was found on the basis of unsupervised PCA of the gene expression data. Previous analysis of methylation groups in CLL suggested a disease-specific role of the transcription factors *EGR, NFAT, AP1* and *EGF* by establishing aberrant methylation patterns^20^. In line with this, we found *NFATC1* and *EGR1* among genes whose expression patterns were associated with methylation groups (Supplemental Figure S6A-B). A detailed analysis of the intermediate methylated subgroup revealed single genes including *SOX11* and *MSI2* that were specific for this subgroup (Supplement Figure S6C-D). *SOX11* is a transcription factor, which expression has been associated with adverse prognostic markers in CLL^36^.

### Expression signature in CLL with trisomy 12

We identified over 3000 differentially expressed genes (with adjusted P-value <0.01) in CLL with trisomy 12 (Figure 1B). Even though chromosome 12 harbours many upregulated genes, the majority of all differentially expressed genes were on other chromosomes and therefore cannot be ascribed to a simple gene dosage effect (Figure 3A).

**Figure 3.**
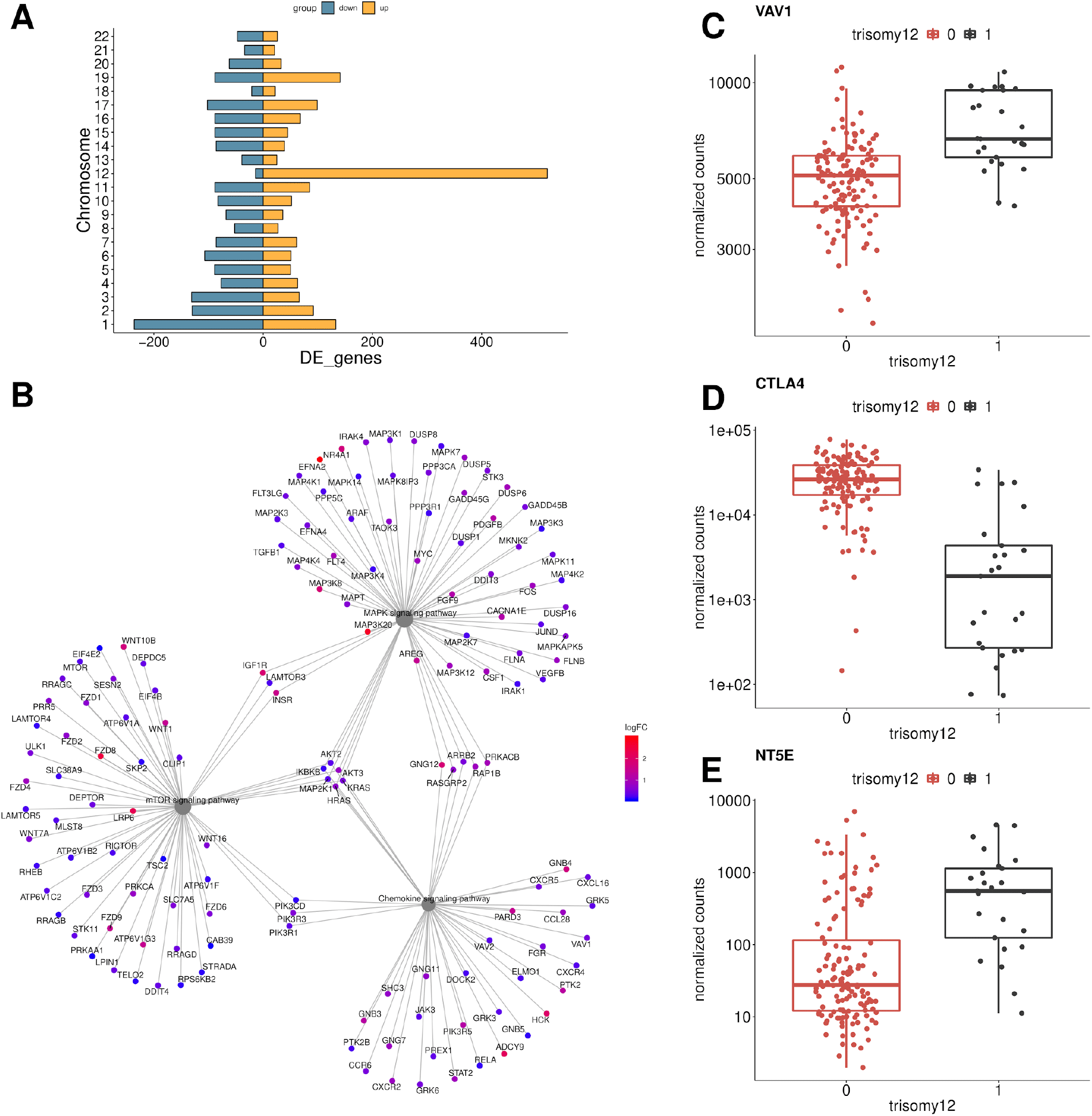
Gene expression in CLL with trisomy 12: A) Role of dosage effect: chromosomal distribution of DE genes in CLL with trisomy 12. Chromosome 12 has the highest number of DE genes, but the majority of DE genes is distributed across all chromosomes and cannot be ascribed to a dosage effect. B) DE genes in enriched KEGG pathways of trisomy 12. C-E) Normalized gene counts of *VAV1, CTLA4* and *NT5E*.

Among the differentially expressed genes, we found numerous genes involved in chemokine signaling such as *VAV1* (Figure 3B,C). Chemokine signaling pathways are induced by chemokine binding and activate MAPK signaling^37,38^. In line with this, we identified differentially expressed genes enriched in MAPK signaling. We also detected an enrichment for the mTOR-signaling pathway, a known modulator of chemokine signaling^39^ (Figure 3B). Consistent with previous reports, integrins like *ITGAL, ITGB2* and *ITGA4* were also upregulated in trisomy 12 samples^13^ (Supplemental table 1). We also found the immune checkpoint gene *CTLA4* (Figure 3D) downregulated in trisomy 12 samples. *CTLA4* has previously been linked to CLL and is associated with increased proliferation and tumor progression^40,41^.

A known mechanism of tumor cells to escape the immune system is to inhibit tumor-specific T cells, and support the conversion of anti-tumor type 1 macrophages to pro-tumor type 2 macrophages by upregulation of ecto-5′-nucleotidase (*NT5E*), which is necessary to convert extracellular ATP into adenosine. A previous study on gene expression in trisomy 12 patients identified *NT5E* as an important element in a trisomy 12 expression network model, but did not directly measure its expression^13^. In our study, we found higher expression of *NT5E* in trisomy 12 and thus can confirm this hypothesis of Abruzzo et al.^13^ (Figure 3E).

Altogether, these results suggest that modulation of MAPK-signaling signaling through chemokine signaling and mTOR-signaling are important mechanisms in trisomy 12 tumorigenesis.

### IGHV status and trisomy 12 affect gene expression in an epistatic way

Epistasis describes a phenomenon where the effect of a genetic variant is dependent on the presence or absence of another genetic variant^42^. Although it is thought to be prevalent, there is almost no data on epistasis between cancer mutations. Because of the major effects of each of these variants individually, we asked whether and to what extent epistatic interactions existed between IGHV mutation status and trisomy 12. We fit a generalized linear model (DESeq2) with main and interaction effects for these covariates. Significant interactions were detected for 893 genes (10% FDR). For these genes, the effect of trisomy 12 was different in U-CLL and M-CLL. We observed four distinct types of epistatic interactions^43,44^ and classified the 893 genes into these categories (Figure 4A-D): *synergy*, where the samples with both variants showed a stronger up-regulation than expected from the single variants; *buffering*, when the presence of both variants led to a strong reduction of gene expression; *inversion*, when the effects in the single variants were reversed in the double variant; *suppression*, when a strong expression change (up or downregulation) of a gene in a single variant was absent in samples with both variants (Figure 4E, G,). Figure 4A-D shows the count data for exemplary genes. Early B-cell factor 1 (*EBF-1*) is up-regulated in all trisomy 12 cases, but this effect is on average about 100 times stronger in M-CLL patients compared to U-CLL patients (synergy) (Figure 4A). While fibroblast growth factor 2 (*FGF2*) is consistently upregulated in trisomy 12 cases with U-CLL, this effect is reversed in M-CLL (suppression) (Figure 4B). Lymphoid enhancer binding factor 1 (*LEF-1*) shows a stable gene expression across samples and has been suggested and tested as a clinical marker for CLL^45^. While the presence of either one of the genetic variants (IGHV-M or trisomy 12) does not seem to have an effect, samples with both of them express consistently lower levels of *LEF1* (buffering) (Figure 4D). These effects cannot be explained by looking at the genotypes independently or by modeling an additive effect of the variants. Each of these interaction types affected dozens or hundreds of genes, and we asked whether there were underlying biological functions. We used gene set enrichment analysis on each of the four types of epistasis, as well as on the combined set of all 893 genes. The overall set of genes with an epistasis expression pattern was enriched in TNF alpha signaling via NF-κB, MYC targets and IL2/STAT5 signaling. A recent study linked NF-κB and EBF1 expression with reduced levels of B-cell signaling^46^. Both IGHV status and trisomy 12 are known to affect BcR signaling. In the type-specific analysis, these pathways were still significantly enriched. In addition, we found the G2M checkpoint pathway enriched in the set of buffered genes, as well as in the set of inversion genes (Figure 4F).

**Figure 4.**
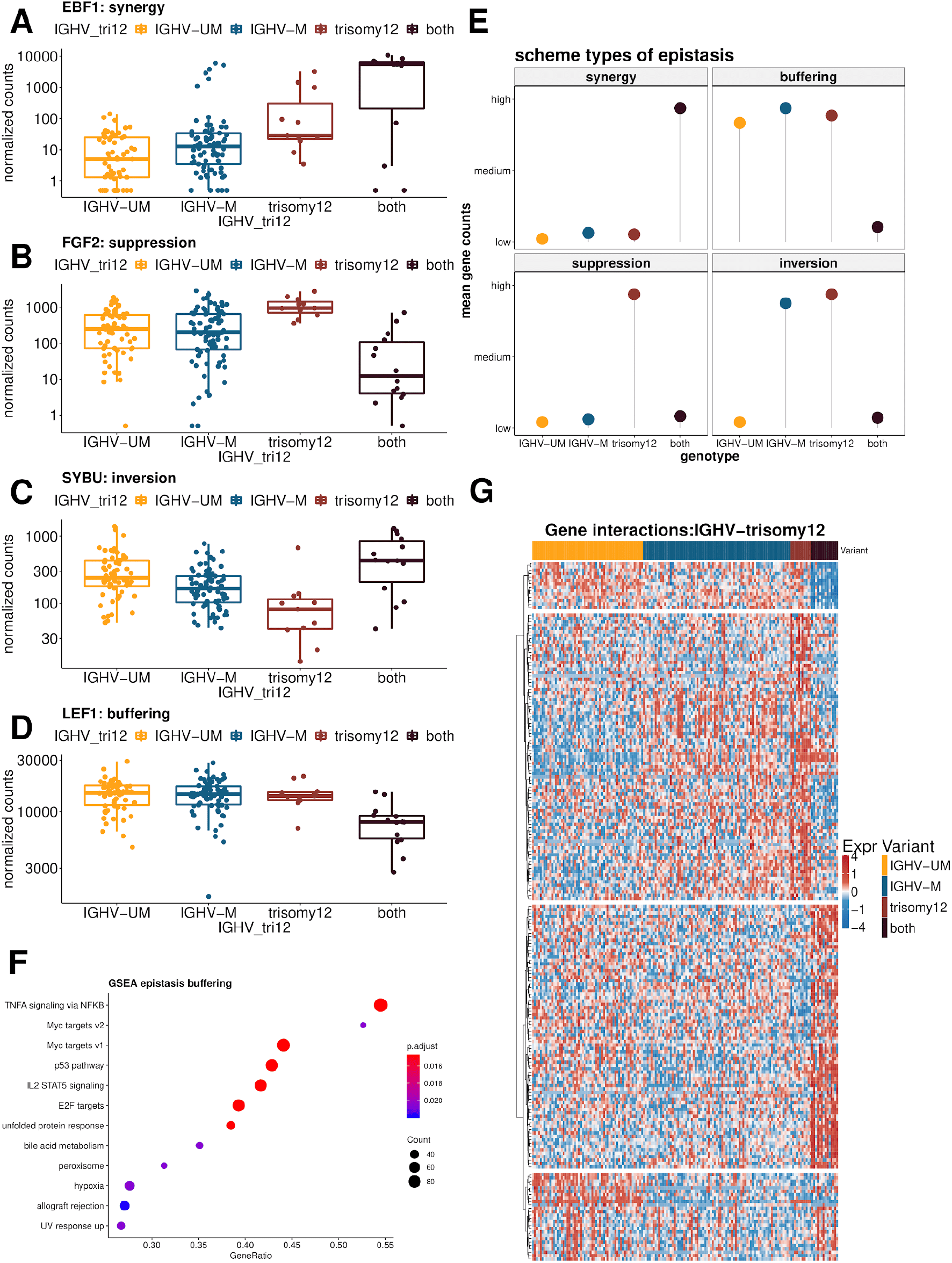
Mixed epistasis of trisomy 12 and IGHV mutation status: A-D) Types of gene expression epistasis: *EBF1* (synergy), *FGF2* (suppression), *SYBU* (inversion), *LEF1* (buffering) E) Schematic classification of epistasis F) Enriched pathways in genes with a buffering epistasis. G) Expression of epistatic gene interactions between trisomy 12 and M-CLL (adjusted P-value < 0.1).

### The epistatic interaction between IGHV status and trisomy 12 affects *ex vivo* drug response in CLL

*Ex vivo* sensitivity to drugs is an informative cellular phenotype that reflects pathway dependencies of CLL cells. We asked whether the epistatic interaction between IGHV mutation status and trisomy 12 on expression level also affected the drug response phenotype. Previously, we measured the *ex vivo* sensitivity, as measured by cell viability, of our 184 CLL samples towards 63 compounds^47^. Again using two-way ANOVA with an interaction term, we identified 6 drugs for which there was a significant (10% FDR) interaction between IGHV mutation status and trisomy 12 (Figure 5). For four drugs, namely, vorinostat, NU7441, fludarabine and AZD7762, we observed a suppression effect, where trisomy 12 led to increased drug sensitivity in the U-CLL, but not the M-CLL samples. For the other two drugs, chaetoglobosin A and BIX02188, the samples with trisomy 12 showed increased resistance particularly in M-CLL, but not in U-CLL.

**Figure 5.**
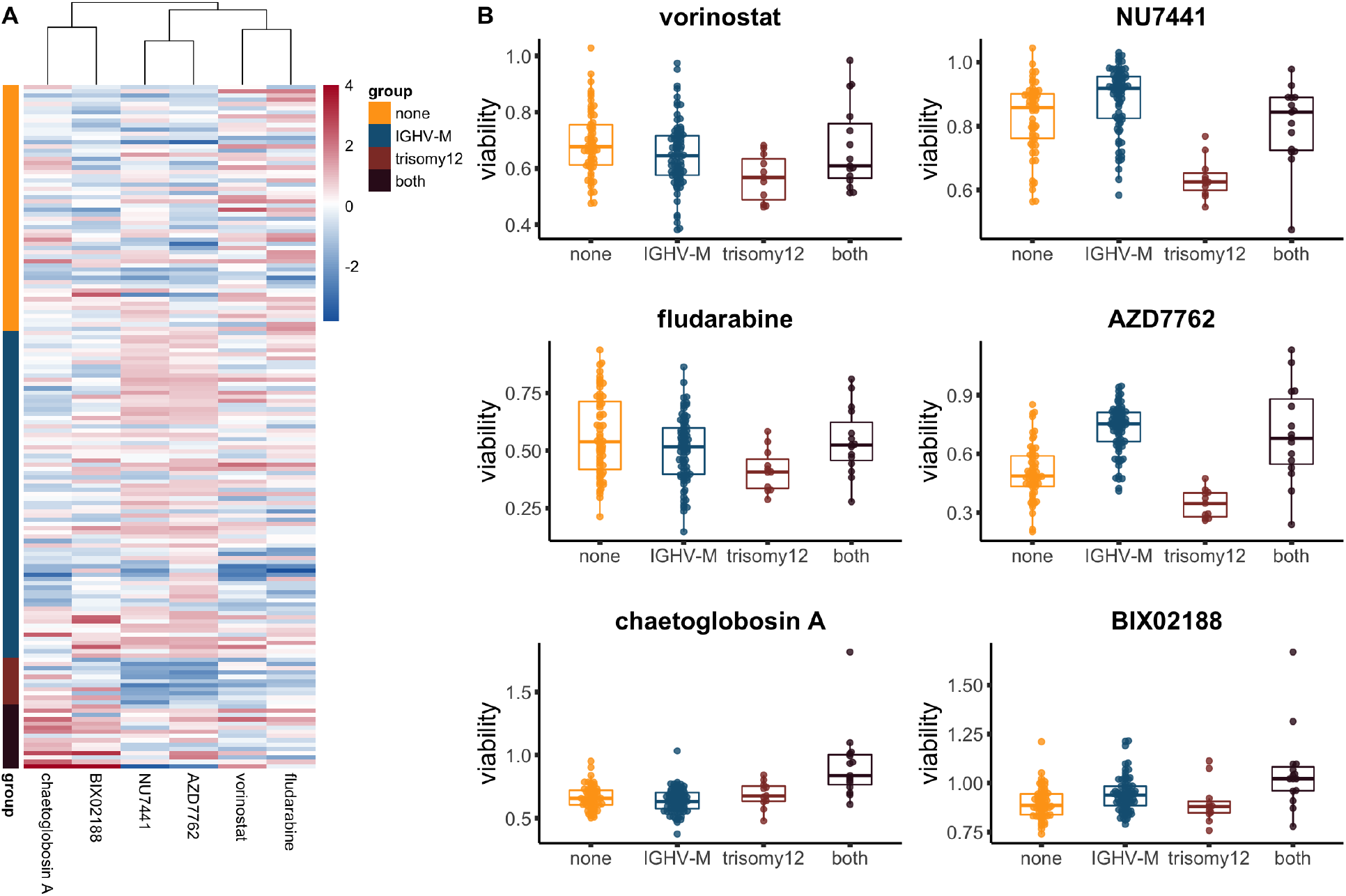
*Ex vivo* drug response phenotype related to the epistatic interaction between IGHV status and trisomy 12, A) Heatmap plot showing the responses of CLL samples (rows) towards the six drugs (columns) for which there was a significant interaction between IGHV mutation status and trisomy 12. The coloring encodes the column-wise z-scores of sample viabilities after drug treatment. B) Boxplots of the viabilities (normalized to DMSO controls) of CLL samples, stratified by their IGHV and trisomy 12 status, towards the six drugs shown in the heatmap.

The four drugs with the suppression phenotype directly or indirectly target DNA: NU7441 inhibits DNA-dependent protein kinase (DNA-PK) and therefore potentiate DNA double-strand breaks^48^; AZD7762 is a checkpoint kinase (CHEK) inhibitor, which can impair DNA repair and increases cell death^49^; fludarabine directly inhibits DNA synthesis and disrupt cell cycle^50^; vorinostat, a histone deacetylase (HDAC) inhibitor, was also reported to induce reactive oxygen species and DNA damage in leukemia cells^51^.

## Discussion

We analyzed 184 CLL transcriptomes, identified gene expression signatures for the most prevalent genetic aberrations and show that these can be used to point towards underlying pathways and potential pathobiological mechanisms. The IGHV mutation status, the three DNA methylation subgroups and trisomy 12 were found as the main drivers of gene expression variability in CLL. This is evident both in unsupervised analyses (clustering, PCA) and in supervised differential expression analysis. Notably, we revealed a much higher impact of IGHV mutation status on the CLL transcriptome than previous reports^10^. The disease stratification by IGHV status is further refined by the three DNA methylation subgroups. We identified genes whose expression follows an apparent continuum from LP, IP to HP in CLL. This finding supports the biological relevance of these three groups and suggests that although they were discovered based on DNA methylation, they are similarly evident at the level of gene expression. Altogether, these results highlight the potential of gene expression profiling to increase our understanding of CLL.

To avoid potential confounding effects of multiple aberrations, other studies have focused on samples with single abnormalities. While this approach could successfully resolve trisomy 12 specific gene expression, it is limited to only a subset of the disease and to selected variants^13^. Here, we demonstrate an improved approach that employs differential expression analysis with generalized linear models and blocking factors, and that is able to use the full range of CLL and to investigate a larger number of genetic aberrations.

Genetic interactions, or epistasis, where the effect of one mutation depends on the genetic status of another locus, is a well known concept in genetics. However, there is surprisingly little data on such phenomena in cancer. Hence, we used the opportunity to study the combinatorial effects of trisomy 12 and the IGHV mutation status on gene expression variability. We identified a large number of genes (∼900) whose expression depended on the presence or absence of either of these two aberrations in a non-additive manner. We categorized these genes into four categories: buffering, synergy, suppression and inversion, each of which contained dozens to hundreds of genes. This means that there is not a single, simple epistasis phenotype between these two aberrations, but a complex, “mixed” pattern. Mixed epistasis of the gene expression phenotypes of pairs of gene alterations has been described in a yeast model system^44^. We employed the genetic interactions between IGHV mutation status and trisomy 12 on gene expression phenotypes to identify pathways, including TNF alpha signaling via NF-κB and the G2M checkpoint pathway, that mediate the effects of these variants. Our results raise the question whether the pathobiology of trisomy 12 may be different between U-CLL and M-CLL, which could be explained by more active BcR signaling in U-CLL and a more diverse set of signaling initiators (independent of the BcR) in M-CLL, and can help us to understand subtype-specific differences.

To our knowledge, this study is the first to provide evidence for this concept in humans, and in cancer. Both the phenomenon itself of epistasis between cancer driving and/or constitutive mutations, and the fact that it can be ‘mixed’ (follow different patterns) for different phenotypes form a new layer of complexity in CLL and in tumor biology more generally. Hence, our study highlights the inherent limitations of studying individual cancer genetic lesions, and points to the need to also map out and understand how they interact and modify each other’s effects.

## Additional Files

Supplementary Table S1: patient_overview.xlsx

Supplementary Table S2: de_genes_all.xlsx

## Acknowledgments

We thank members of the Huber and Zenz research teams for valuable discussions. The work was supported by the European Union (Horizon 2020 project SOUND under grant agreement number 633974) and the German Federal Ministry of Education and Research (TRANSCAN project GCH-CLL 143 under grant agreement number 01KT1610 and CompLS project MOFA under grant agreement number 031L0171A). T. Zenz was supported by the UZH Clinical Research Priority Program “Next Generation Drug Response Profiling for Personalized Cancer Care “, the Swiss Cancer League (KFS-4439-02-2018), and the Monique-Dornonville-de-la-Cour Stiftung. R. Rosenquist received funding from the Swedish Cancer Society, the Swedish Research Council, the Knut and Alice Wallenberg Foundation, and Radiumhemmets Forskningsfonder, Stockholm. For technical support and expertise, we thank the DKFZ Genomics and Proteomics Core Facility. We thank Hanno Glimm, Stefan Fröhling, Daniela Richter, Roland Eils, Peter Lichter, Stephan Wolf, Katja Beck, and Janna Kirchhof for infrastructure and program development within DKFZ-HIPO and NCT POP.

## Authorship Contributions

A Lütge: conceptualization, data curation, formal analysis, visualization, methodology, and writing - original draft, review, and editing.

J. Lu: conceptualization, formal analysis, methodology, and writing - original draft, review, and editing.

J. Hüllein: data curation

T. Walther: data curation

L. Sellner: data curation, - review, and editing

B. Wu: data curation, - review, and editing

R. Rosenquist: data curation, writing - review, and editing

C. Oakes: data curation, - review, and editing

S. Dietrich: data curation, - review, and editing

W. Huber: conceptualization, supervision, methodology, project administration, and writing - original draft, review, and editing.

T. Zenz: conceptualization, supervision, methodology, project administration, and writing - original draft, review, and editing.

