## Supplemental Figures for "Subgroup-specific gene expression profiles and mixed epistasis in chronic lymphocytic leukemia"

### Supplement

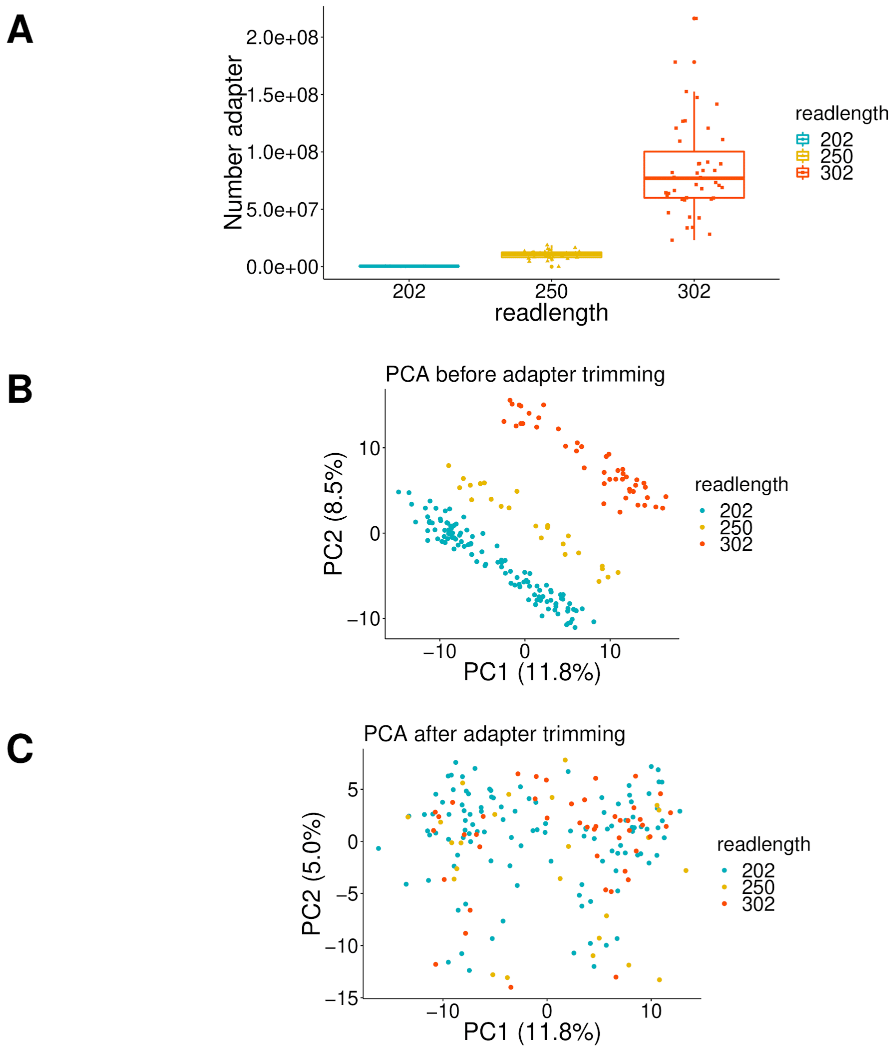


**Supplemental Figure S1. Effect of adapter trimming on sequencing batches,** A) The number of reads with a part of their sequence mapping to the adapter sequences increases by read length. B) PC1 and PC2 are related to batch differences due to differences in read length, but not C) after adapter trimming.

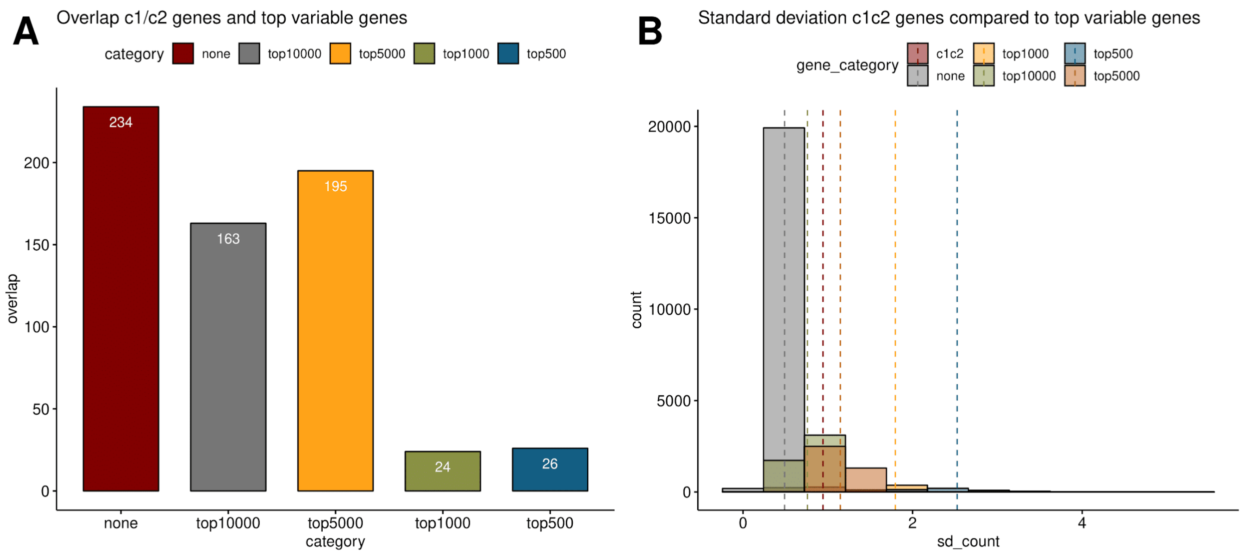


**Supplemental Figure S2. DE genes of c1/c2 groups as observed by Ferreira et al.[10],** A) Overlap of DE genes between c1/c2 groups and the 10000 resp. 5000, 1000 and 500 most variable genes. Only 51 DE genes between c1/c2 groups are in the 1000 most variable genes. B) Standard deviation of most variable genes compared to c1/c2 genes.

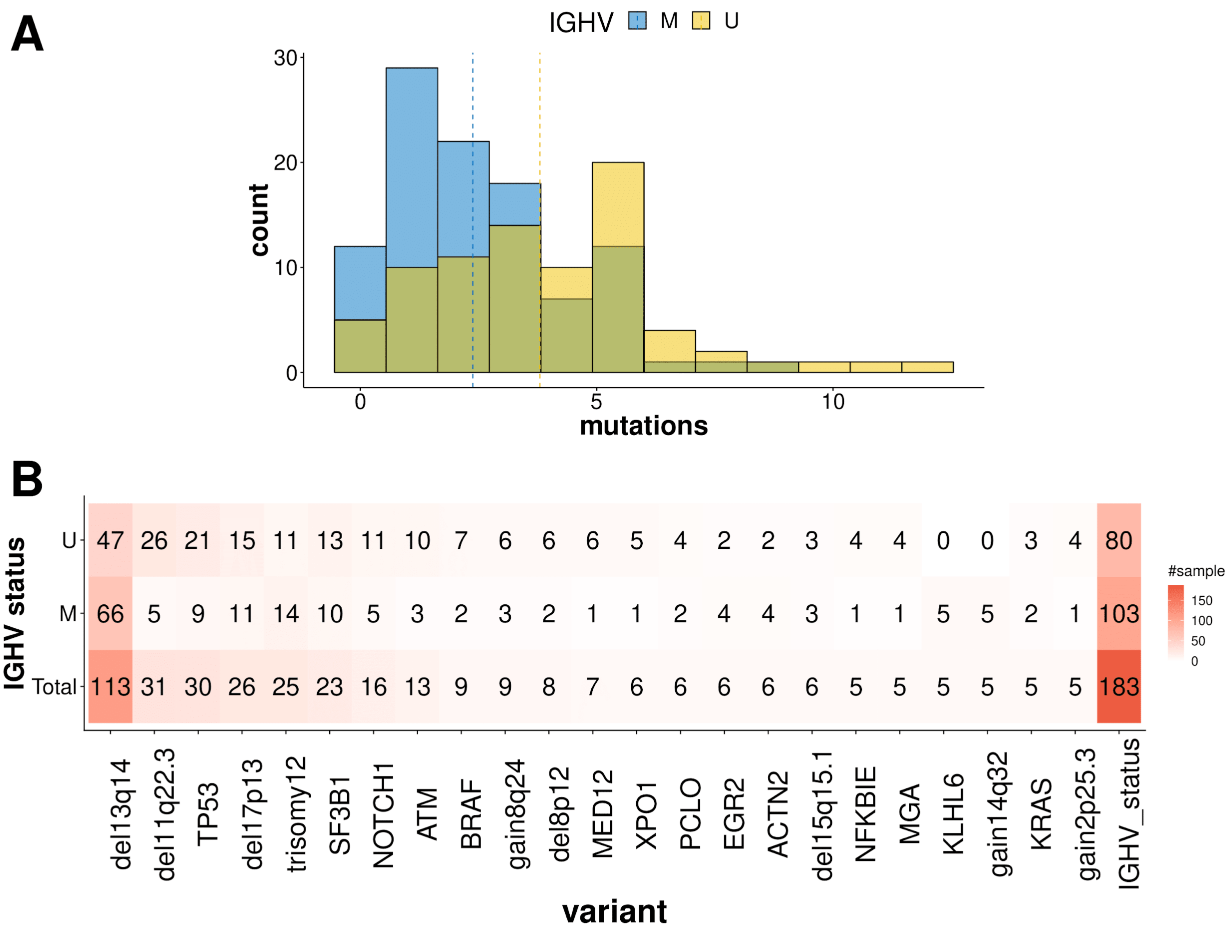


**Supplemental Figure S3. Mutational load by sample,** A) The number of mutations (including genetic variations) by sample. On average M-CLL samples have 2.6 and U-CLLsamples 4 genetic aberrations. B) Number of samples per genetic variant explored included in this study by IGHV status.

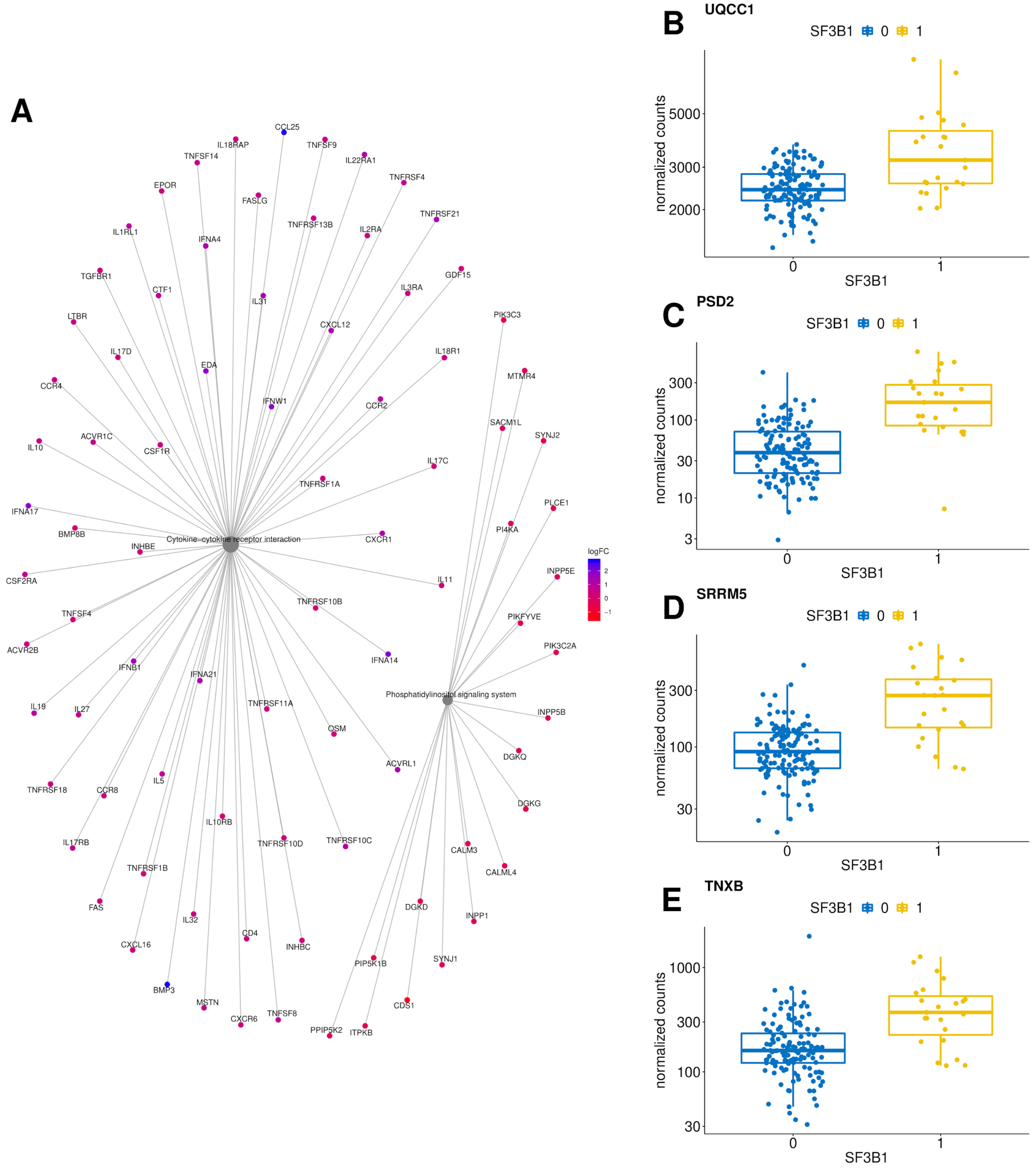


**Supplemental Figure S4. Gene expression associated with *SF3B1*:** A) Differentially expressed genes in enriched KEGG pathways of SF3B1. B-E) Normalized gene counts of *UQCC1*, *PSD2*, *SRRM5* and *TNXB*.

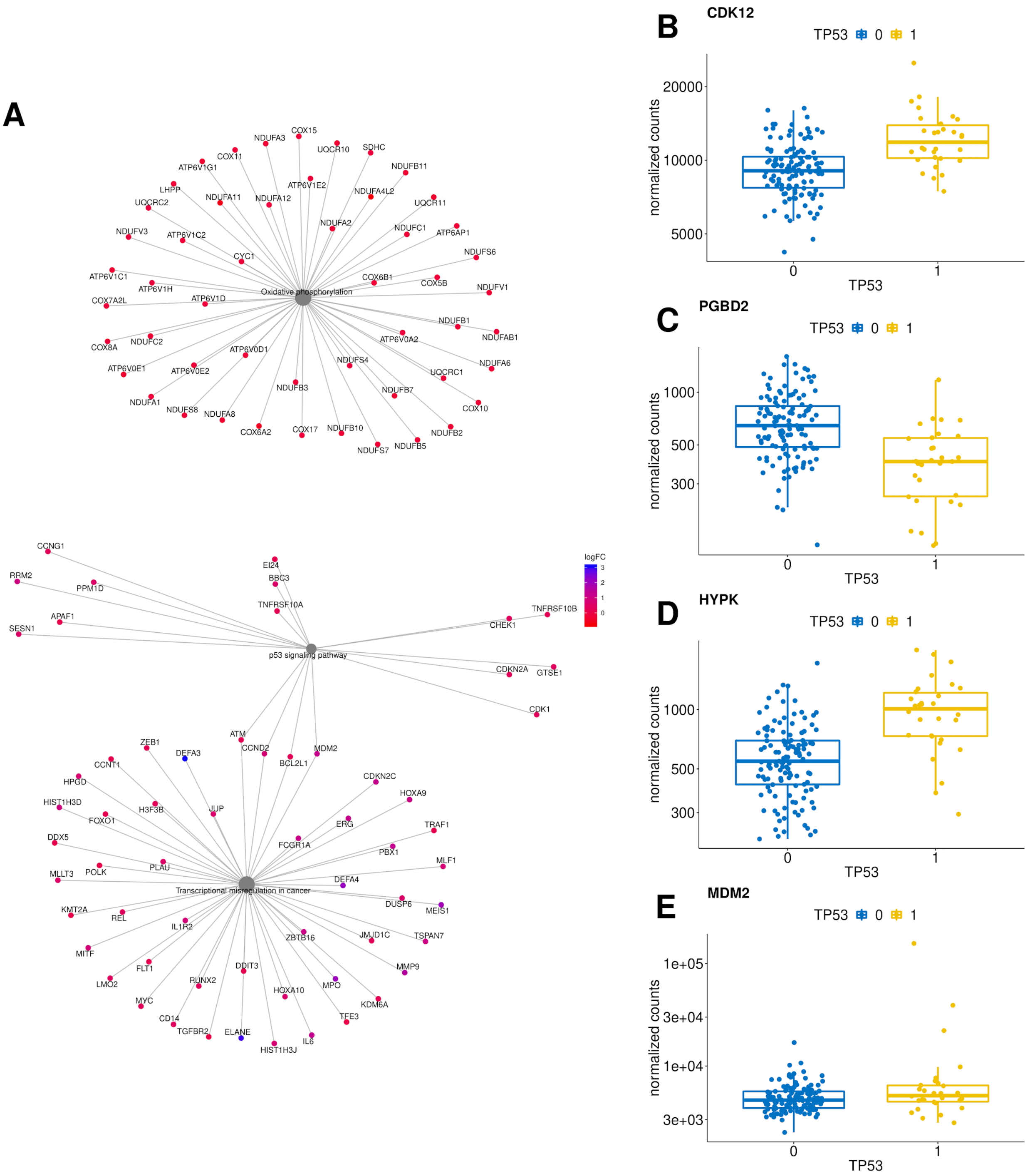


**Supplemental Figure S5: Gene expression associated with *TP53*:** A) Differentially expressed genes in enriched KEGG pathways of TP53. B-E) Normalized gene counts of *CDK12*, *PGBD2*, *HYPK* and *MDM2*.

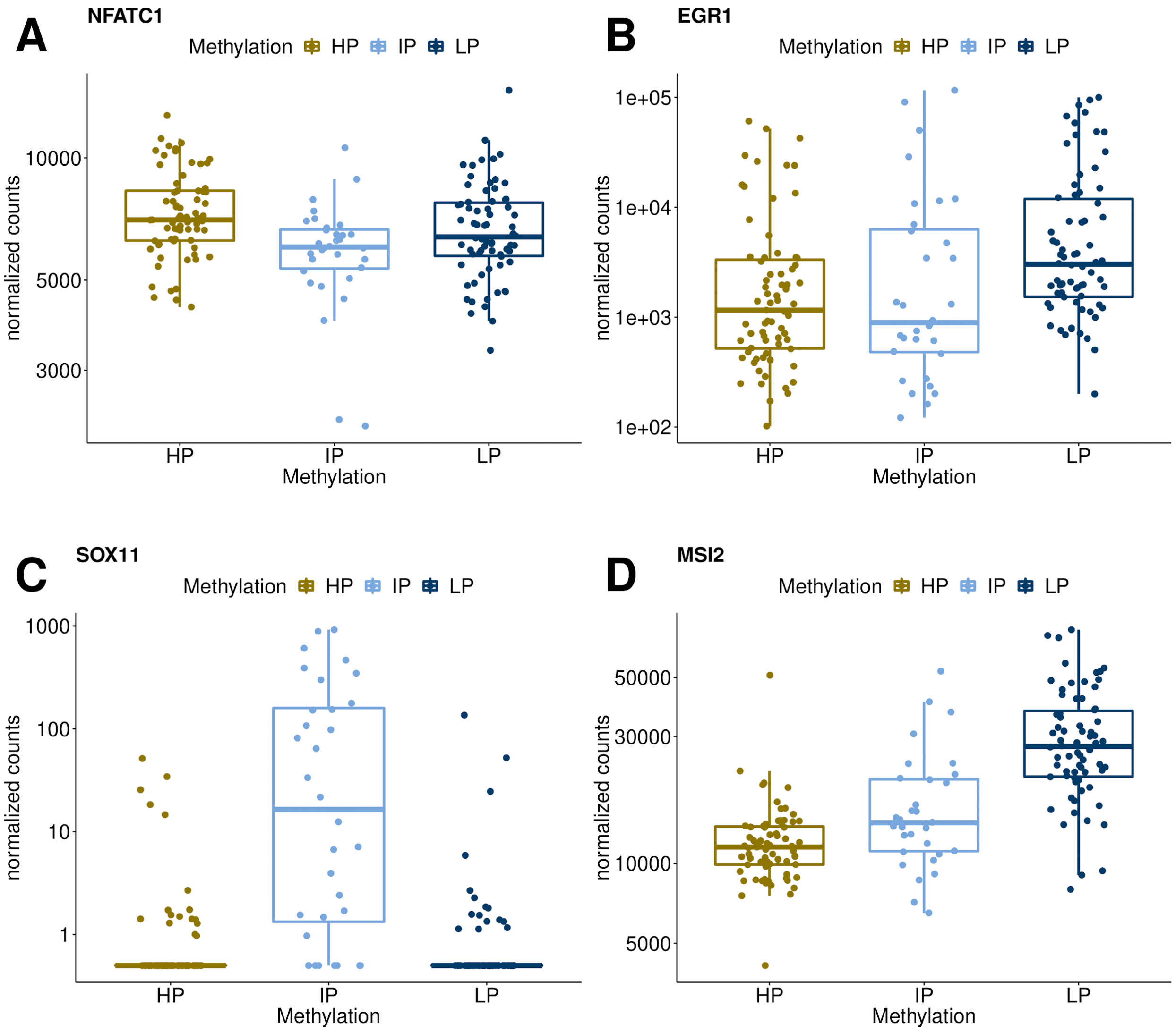


**Supplemental Figure S6. Gene expression associated with HP, IP and LP groups:** A-D) Normalized gene counts of *NFATC1*, *EGR1*, *SOX11* and *MSI2*.
